# Ultrastructural insight into SARS-CoV-2 attachment, entry and budding in human airway epithelium

**DOI:** 10.1101/2021.04.10.439279

**Authors:** Andreia L Pinto, Ranjit K Rai, Jonathan C Brown, Paul Griffin, James R Edgar, Anand Shah, Aran Singanayagam, Claire Hogg, Wendy S Barclay, Clare E Futter, Thomas Burgoyne

## Abstract

Ultrastructural studies of SARS-CoV-2 infected cells are crucial to better understand the mechanisms of viral entry and budding within host cells. Many studies are limited by the lack of access to appropriate cellular models. As the airway epithelium is the primary site of infection it is essential to study SARS-CoV-2 infection of these cells. Here, we examined human airway epithelium, grown as highly differentiated air-liquid interface cultures and infected with three different isolates of SARS-CoV-2 including the B.1.1.7 variant (Variant of Concern 202012/01) by transmission electron microscopy and tomography. For all isolates, the virus infected ciliated but not goblet epithelial cells. Two key SARS-CoV-2 entry molecules, ACE2 and TMPRSS2, were found to be localised to the plasma membrane including microvilli but excluded from cilia. Consistent with these observations, extracellular virions were frequently seen associated with microvilli and the apical plasma membrane but rarely with ciliary membranes. Profiles indicative of viral fusion at the apical plasma membrane demonstrate that the plasma membrane is one site of entry where direct fusion releasing the nucleoprotein-encapsidated genome occurs. Intact intracellular virions were found within ciliated cells in compartments with a single membrane bearing S glycoprotein. Profiles strongly suggesting viral budding from the membrane was observed in these compartments and this may explain how virions gain their S glycoprotein containing envelope.

## Introduction

Severe acute respiratory syndrome coronavirus-2 (SARS-CoV-2) emerged as a novel virus in late 2019, causing widespread infections within a short period of time leading to a global pandemic. The high affinity of SARS-CoV-2 Spike (S) glycoprotein for human angiotensin-converting enzyme 2 (ACE2) receptor has been proposed as the likely cause of rapid spread of infection (Shang et al., 2020). ACE2 is expressed at the apical surface of primary airway epithelial cells, with expression levels showing a gradient from upper to lower airway (Hou et al., 2020). S glycoprotein forms homotrimers emanating from the viral surface and comprises the S1 subunit with a receptor-binding domain (RBD) that binds to ACE2 and the S2 subunit that mediates fusion of viral and cellular membranes (Yang et al., 2020).

The serine protease TMPRSS2 that is also expressed within the respiratory epithelium has been implicated in viral entry by cleaving S glycoprotein (Hoffmann et al., 2020). Upon binding to ACE2, the prefusion S1/S2 complex is proteolytically cleaved at a second site in S2 (S2’) by TMPRSS2, which triggers dissociation of S1 and a conformational shift in S2 to give the post-fusion form of S glycoprotein (Cai et al., 2020). Fusion of viral membrane to the host cell plasma membrane has been shown to be driven by this cleaved membrane anchored S2 subunit (Walls et al., 2020). After membrane fusion occurs, the nucleocapsid protein complexed with viral RNA are released from the viral lumen into the cytoplasm of the host cell to initiate viral replication.

SARS-CoV-2 is able to enter cells lacking TMPRSS2 by alternative pathways (Sungnak et al., 2020; Hoffmann et al., 2020; Peacock et al., 2020). Endocytosed virus is trafficked to endosomes that eventually fuse with lysosomes and/or are engulfed by autophagosomes (Yang and Shen, 2020). Endocytic cysteine proteases cathepsins B and L can cleave the S2’ site to gain a postfusion equivalent form of S glycoprotein (Millet and Whittaker, 2014; Cai et al., 2020). There is evidence that two-pore channels can facilitate the release of viral RNA from endolysosomes to allow for viral replication (Khan et al., 2020).

Virus replication is believed to take place at perinuclear sites where like for other positive-sense RNA viruses, endoplasmic reticulum (ER)-derived structures known as double-membrane vesicles (DMVs) form (Snijder et al., 2020). The DMVs are interconnected by convoluted membranes and small open double-membrane spherules (DMSs) (V’kovski et al., 2021). DMVs are enriched in double-stranded RNA and are associated with viral replication (Klein et al., 2020). SARS-CoV-2 viral budding occurs at ER-to-Golgi intermediate compartments (ERGIC) as detected in SARS-CoV (Stertz et al., 2007). It has also been suggested that the virus can exploit lysosomes to allow egress out of the host cell post viral replication (Ghosh et al., 2020). The precise route from DMV to viral budding to egress of budded virions from the cell remains to be fully established. Cleavage at the S1/S2 junction may occur during trafficking by host furin-like enzymes before reaching the cell surface (Matsuyama et al., 2010). There is evidence this pre-cleavage of spike enhances entry into respiratory tissue and that variants that lack the furin cleavage site use the endosomal pathway for entry and are inefficiently transmitted by the respiratory route (Peacock et al., 2020).

Novel SARS-CoV-2 variants have emerged with genetic constellations that include mutations in S glycoprotein. This raises concern as there is potential for differences in transmissibility rates, severity of disease and resistance to neutralizing antibodies. Lineage B.1.1.7 (Variant of Concern 202012/01) is a variant that emerged in the UK in September 2020 (Rambaut et al., 2020). By early 2021, B.1.1.7 has become the dominant lineage in the UK, likely due to its increased transmissibility (Volz et al., 2021). 23 mutations have been identified across the viral genome, including 9 associated with S glycoprotein including N501Y in RBD, Δ69-70, Δ144 in NTD and P681H close to the furin cleavage site (Rambaut et al., 2020; Brown et al., 2021). Contemporaneous lineages which do not demonstrate the apparent increased transmissibility of B.1.1.7 include B.1.258 that has Δ69-70 and N439K mutations and B.1.117.19 with a A222V mutation in S glycoprotein (Brown et al., 2021).

In this study, we examined the ultrastructure of human respiratory epithelial cells infected with three different isolates of SARS-CoV-2 (B.1.1.7, B.1.258 and B.1.117.19). No differences in the characteristics of infection were observed but the similarities provide insight into viral attachment, entry and budding in human airway epithelium (HAE).

## Methods

### Human nasal brushing biopsies

Nasal brushings samples were taken from healthy and SARS-CoV-2 infected participant turbinates using 3-mm bronchial cytology brushes under Health Research Authority study approval (REC ref: 20/SC/0208; IRAS: 282739). SARS-CoV-2 infection was PCR-confirmed from swabs taken at the Royal Brompton and Harefield Hospital.

### Human airway epithelium (HAE) cell culture

Nasal brushings were placed in PneumaCult-Ex Plus medium (STEMCELL Technologies, Cambridge, UK) and cells dissociated from the brush by gentle agitation. The cells were seeded into a single well of a collagen (PureCol from Sigma Aldrich) coated plate and once confluent, the cells were passaged and expanded further in a T25 flask. The cells were passaged a second time and seeded onto transwell inserts (6.5 mm diameter, 0.4 μm pore size, Corning) at a density of 24,000 cells per insert. Cells were cultured in PneumaCult-Ex Plus medium (STEMCELL Technologies, Cambridge, UK) until confluent, at which point the media was replaced with PneumaCult-ALI medium in the basal chamber and the apical surface exposed to provide an air liquid interface (ALI). Ciliation was observed between 4-6 weeks post transition to ALI.

### HeLa cell culture and transfections

HeLa cells were seeded onto glass coverslips and transfected with either myc-ACE2 (pCEP4-myc-ACE2 from Addgene Plasmid #141185), ACE2 (Daly et al., 2020), HA-TMPRSS2 (pCSDest-HA-TMPRSS2 from Addgene Plasmid #154963), ss-HA-Spike plasmids (HA tag at the N terminus with a Serine-Glycine linker between residues S13 and Q14 of S glycoprotein (Stewart et al., 2021)) were transfected using Lipofectamine 3000 (ThermoFisher Scientific), following the manufacturers guidelines. Cells were fixed for immunofluorescence or electron microscopy 2 days after transfection.

### Viruses and infection of HAE cells

SARS-CoV-2 B.1.1.7 isolate hCoV-19/England/204661721/2020 (EPI_ISL_693400), B.1.258 isolate hCoV-19/England/204501206/2020 (EPI_ISL_660791) and B.1.117.19 isolate hCoV-19/England/204501194/2020 (EPI_ISL_660788) were isolated from swabs as described in (Brown et al., 2021). Swabs were collected by the PHE Virology Consortium and ATACCC under the Integrated Network for Surveillance, Trials and Investigation of COVID-19 Transmission (INSTINCT; Ethics Ref: 20/NW/0231; IRAS Project ID: 282820) The investigation protocol was reviewed and approved by the PHE Research Ethics and Governance Group and Incident Management team. PHE has legal permission, provided by Regulation 3 of the Health Service (Control of Patient Information) Regulation 2002, to process patient confidential information for national surveillance of communicable diseases. Isolates were passaged twice in Vero cells before being used to infect HAE cells. To remove the mucus layer from the apical surface of HAE cells prior to infection, 200ul of DMEM was added and incubated at HAE cells were infected at a 37°C, 5% CO_2_ for 10 mins before removal of medium. HAE cells were infected at a multiplicity of infection (MOI) of 0.01 pfu/cell of each isolate diluted in DMEM. Inocula were added to the apical chamber and incubated for 1 h at 37°C, 5% CO_2_ before removal of the inoculum and incubating for a further 72 hr. Subsequently, cells were fixed for electron microscopy.

### Conventional transmission electron microscopy (TEM)

Cultured HAE cells, nasal brushing samples and HeLa cells were fixed by placing them in 2.5% glutaraldehyde in 0.05M sodium cacodylate buffer at a pH 7.4 and left for at least 24 hours. Subsequently, the samples were incubated in 1% aqueous osmium tetroxide for 1 h at room temperature (RT) before en bloc staining in undiluted UA-Zero (Agar Scientific) for 30 minutes at RT. The samples were dehydrated using increasing an ethanol series (50%, 70%, 90%, 100%), followed by propylene oxide and a mixture of propylene oxide and araldite resin (1:1). The samples were embedded by placing them in araldite and left at 60°C for 48 h. Ultrathin sections were cut using a Reichert Ultracut E ultramicrotome and stained using Reynold’s lead citrate for 10 minutes at RT. Images were acquired on a JEOL 1400Plus TEM fitted with an Advanced Microscopy Technologies (AMT) XR16 charge coupled device (CCD) camera.

### Electron tomography (ET)

Electron microscopy sections were tilted and imaged within a JEOL 1400Plus TEM from ±60° over two perpendicular axes and dual axis tomograms were generated using IMOD (Kremer et al., 1996). Subtomographic averaging was performed using the IMOD PEET (Particle Estimation for Electron Tomography) package. Slices from the tomograms were viewed in IMOD and the averaged structures surface rendered in Chimera (UCSF).

### Immunofluorescence (IF)

HeLa and HAE cells as well as nasal brushing biopsies were fixed in 4% PFA in PBS for 1 hour. HeLa cells were stained directly whereas HAE and nasal brushing were embedded and frozen in OCT (optimal cutting temperature) compound before cutting ~15 μm thick sections using a cryostat. HeLa cells and sections were permeabilised using 0.2% saponin in PBS for 20 mins at room temperature before incubating in blocking solution containing 0.02%, 1% BSA in PBS for 30 mins at room temperature. HeLa cells on coverslips were incubated with primary antibodies for 1 hr at room temperature whereas cryostat sections were incubated overnight at 4°C. Antibodies against ACE2 (Abcam and Sigma-Aldrich), TMPRSS2 (Novus Biologicals), HA (Biolegend) and Myc (Santa Cruz), Ezrin (Santa Cruz) were used. Alex Fluor bound secondary antibodies (ThermoFisher Scientific) and phalloidin 647 (Abcam) were applied to samples for 1 hour at room temperature. Images were acquired using a Leica SP8 confocal and analysed using ImageJ.

## Results

### SARS-CoV-2 isolates show similar characteristics upon infection of human airway cells

The nose is the primary site of SARS-CoV-2 infection and therefore nasal primary human airway epithelial (HAE) cells are the optimal cell culture model to simulate SARS-CoV-2 infection of the airway. Differentiation of cells at an air liquid interface (ALI) results in an epithelial layer composed of ciliated, goblet and basal cells. We previously infected HAE cells with a panel of SARS-CoV-2 isolates at an MOI of 0.01 pfu/cell and demonstrated robust viral replication kinetics in these cells with abundant infectious virus released at the apical surface at 72 hours post infection (Brown et al., 2021). Fixation of these samples at 72 hours post-infection and observation by EM showed the presence of SARS-CoV-2 virions and major changes in cellular morphology as shown in Figure 1. As described in previous studies, host cell membrane remodelling to give way to viral replication machinery was observed, which included DMVs, convoluted membranes (CMs) and viral containing compartments (VCs) (Figure 1, C – D) (Snijder et al., 2020; Zhu et al., 2020). Some DMVs were seen to be connected to VCs as shown by the blue arrowhead in Figure 1 E. At the 72 hour post infection time point, substantial numbers of virions visible in the extracellular space were attached to the plasma membrane (red arrowheads Figure 1 D, F) consistent with the high viral titres observed in washings of the cells at this time (Brown et al., 2021). Given the initial low MOI with which the cells were treated this must have been virus resulting from replication and release from infected cells potentially attaching to initiate further rounds of infection.

**Figure 1.**
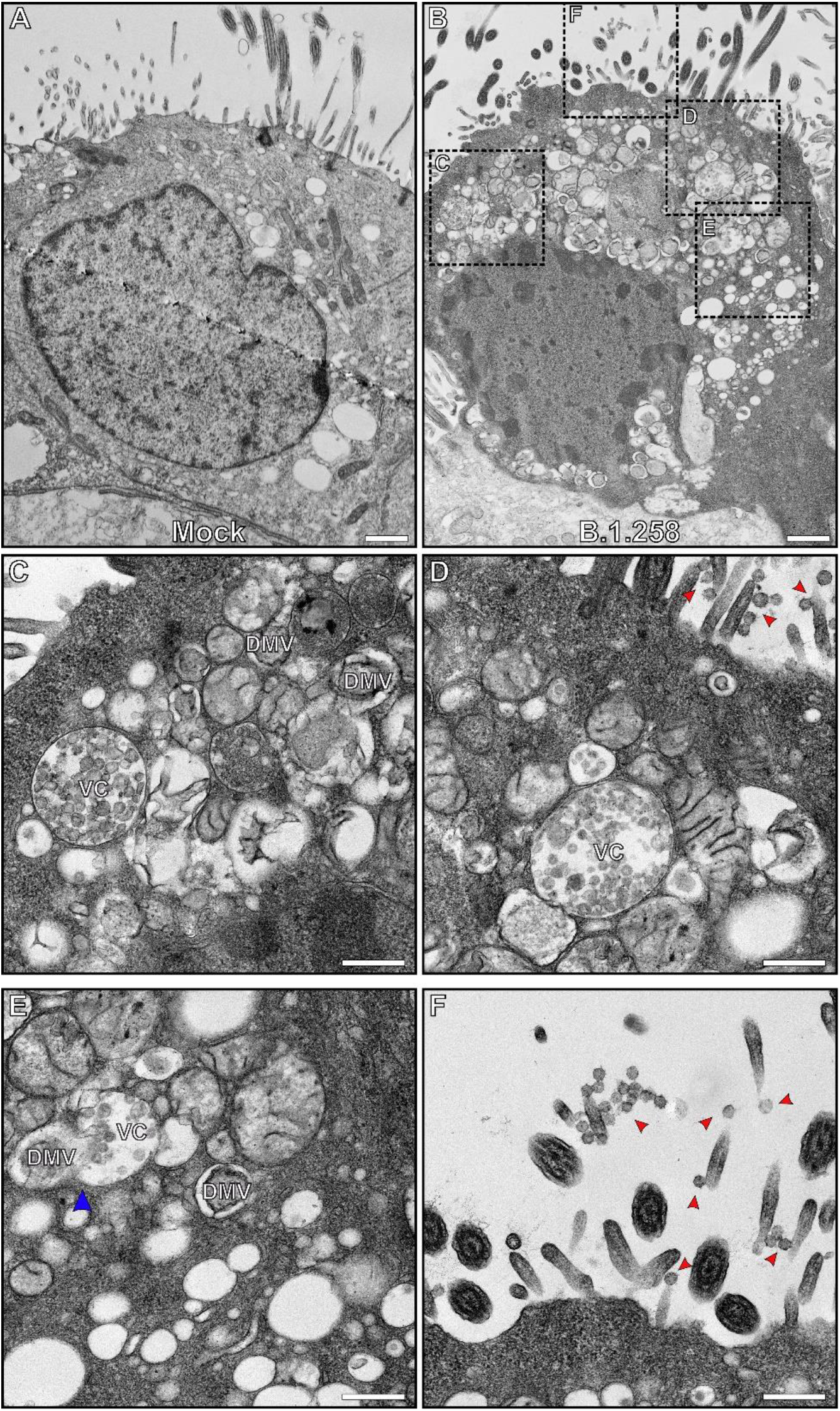
Human nasal respiratory epithelium infected with SARS-CoV-2 show considerable remodelling to give way to viral replication. (A) A non-infected ciliated epithelial cell. (B) A cell infected with B.1.258 SARS-CoV-2 variant contains a large number of viral associated compartments shown at higher magnification in (C – E) and virions at the cell surface (red arrowheads) (D, F). (C – E) Viral containing compartments (VCs) and double membrane vesicles (DMVs) can be seen. (E) Some DMVs are connect to VCs (blue arrowhead) and there are convoluted membrane surrounding these compartments. Scalebars (A & B) 1 μm and (C – F) 400 nm.

One of the identifiable traits of SARS-CoV-2 virions by TEM is the nucleocapsid contents of the viral lumen that present as an electron dense punctate pattern, making them distinguishable from clathrin coated and intraluminal vesicles (Figure 2). The SARS-CoV-2 spike (S) glycoprotein was evident on most extracellular virions as well as those in VCs (Figure 2 A). Some of the VCs contained virions that were not coated with S glycoprotein on the surface. When comparing the three isolates B.1.1.7, B.1.258, B.1.117.19, at the resolution of conventional TEM there were no obvious differences in structure (Figure 2B).

**Figure 2.**
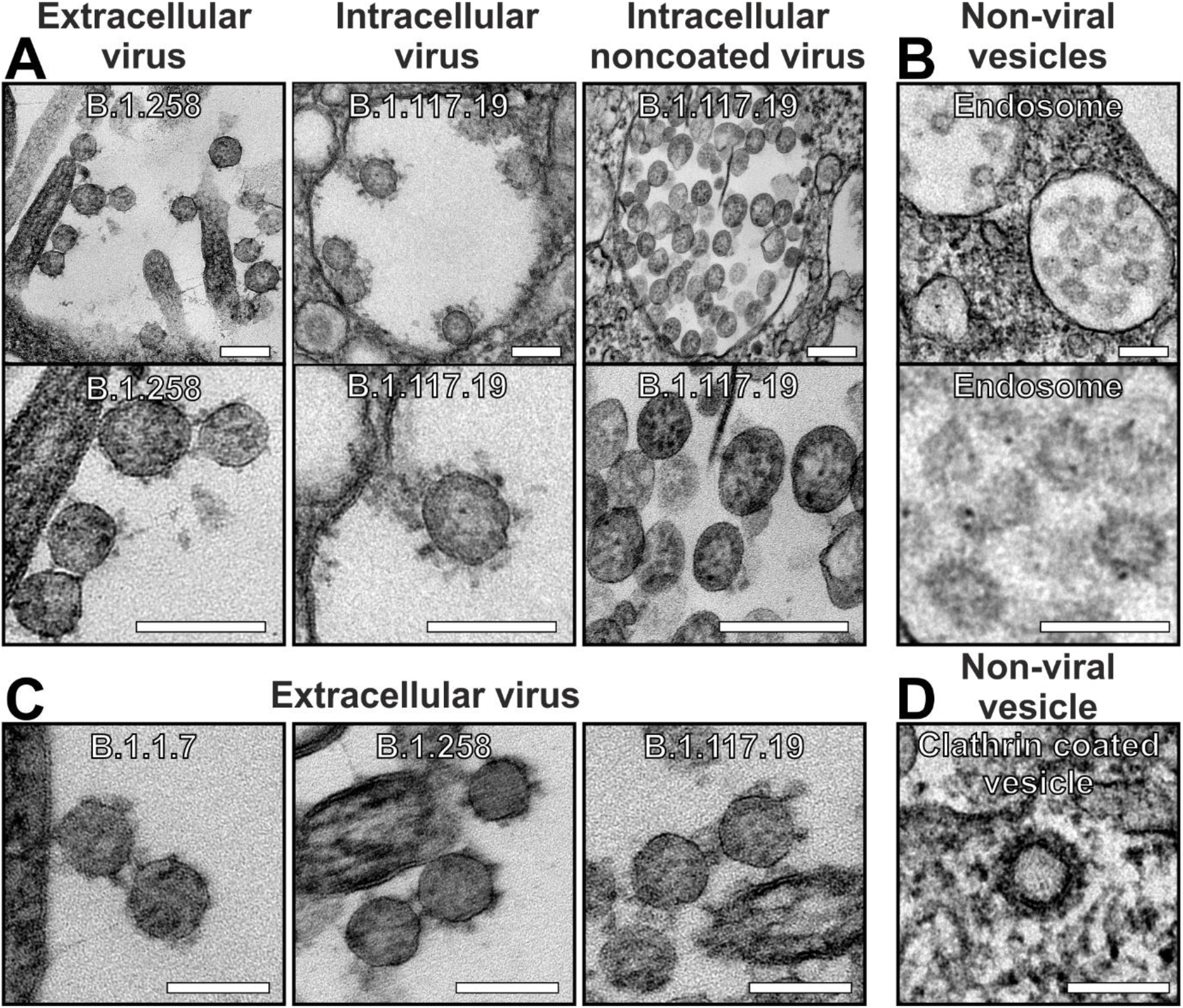
By TEM, SARS-CoV-2 virions are discernible by the punctate pattern of the nucleocapsid containing lumen and often the presence of a S-glycoprotein coat. (A) Virions are seen extracellularly at the cell surface and intracellularly within viral containing compartments. In some compartments virions were observed without a S-glycoprotein coat. (B) In comparison, endosomes consist of intraluminal vesicles that are smaller, lighter in appearance and lack the nucleocapsid punctate interior of virions. (C) When comparing three different lineages (B.1.1.7, B.1.258 and B.1.117.19) there were no obvious differences in structure. (D) A clathrin coated vesicle can be discerned from a virion by having a denser coat that viral S-glycoprotein and lacking punctate nucleocapsid in the interior. Scalebars (A - D) 100 nm.

### Extracellular virions do not localise to ciliary membranes

Sections of HAE were stained for the extracellular and intracellular domains of ACE2 and the intracellular domain of TMPRSS2 by immunofluorescence. This was done in combination with antibody staining against tubulin to identify cilia and phalloidin to highlight the actin enriched microvilli at the cell surface. The staining shows ACE2 and TMPRSS2 localised to regions of plasma membrane including microvilli but excluded from cilia (Figure 3 A – D). Antibody detecting the intracellular domain of ACE2 stained intensely at the base of microvilli, whereas staining with antibody against the extracellular ACE2 domain also extended through the microvilli level. Neither antibody stained the tips of cilia. The difference in staining pattern between the two ACE2 antibodies could result from the extracellular domain antibody only detecting the larger ACE2 isoform, whereas the intracellular domain antibody is detecting the short ACE2 isoform (Blume et al., 2021). The shorter isoform may not be localised through the microvilli, therefore, detection of this and the longer isoform of ACE2 at the microvilli base by the intracellular domain antibody would generate a stronger signal that is more readily detectable compared to the longer ACE2 isoform through the microvilli. A similar staining pattern was seen in a human nasal brushing biopsy showing that the protein localisation is not an artefact of cell culture (Supplementary Figure 1). To validate the ACE2 and TMPRSS2 antibodies that were used, they were tested in HeLa cells expressing ACE2, Myc-ACE2 or HA-TMPRSS2 (Supplementary Figure 2), in addition to the extracellular ACE2 antibody applied to non-permeabilised sections (Supplementary Figure 3). When looking at non-permeabilised sections there was diffuse tubulin staining due to poor accessibility of the antibody into cilia, but the antibody raised against the extracellular domain of ACE2 still provided a strong signal. This was particularly evident in a confocal slice of apical epithelium where cilia were absent and ACE2 was seen associated with the microvilli (phalloidin stained) (Supplementary Figure 3). When using an antibody against Ezrin, an actin binding component of the microvilli, the staining surrounded the actin (phalloidin stained) similar to the antibody against the extracellular domain of ACE2 (Figure 3 B and Supplementary figure 3 & 4). Consistent with the localisation of key entry molecules on the plasma membrane at the base of and surrounding the microvilli but not on the cilia, TEM showed extracellular virions at the surface of ciliated epithelial cells mostly in contact with microvilli but not cilia (Figure 3 E-G). The preferential attachment to microvilli over cilia was observed for all three isolates (Figure 3 H).

**Figure 3.**
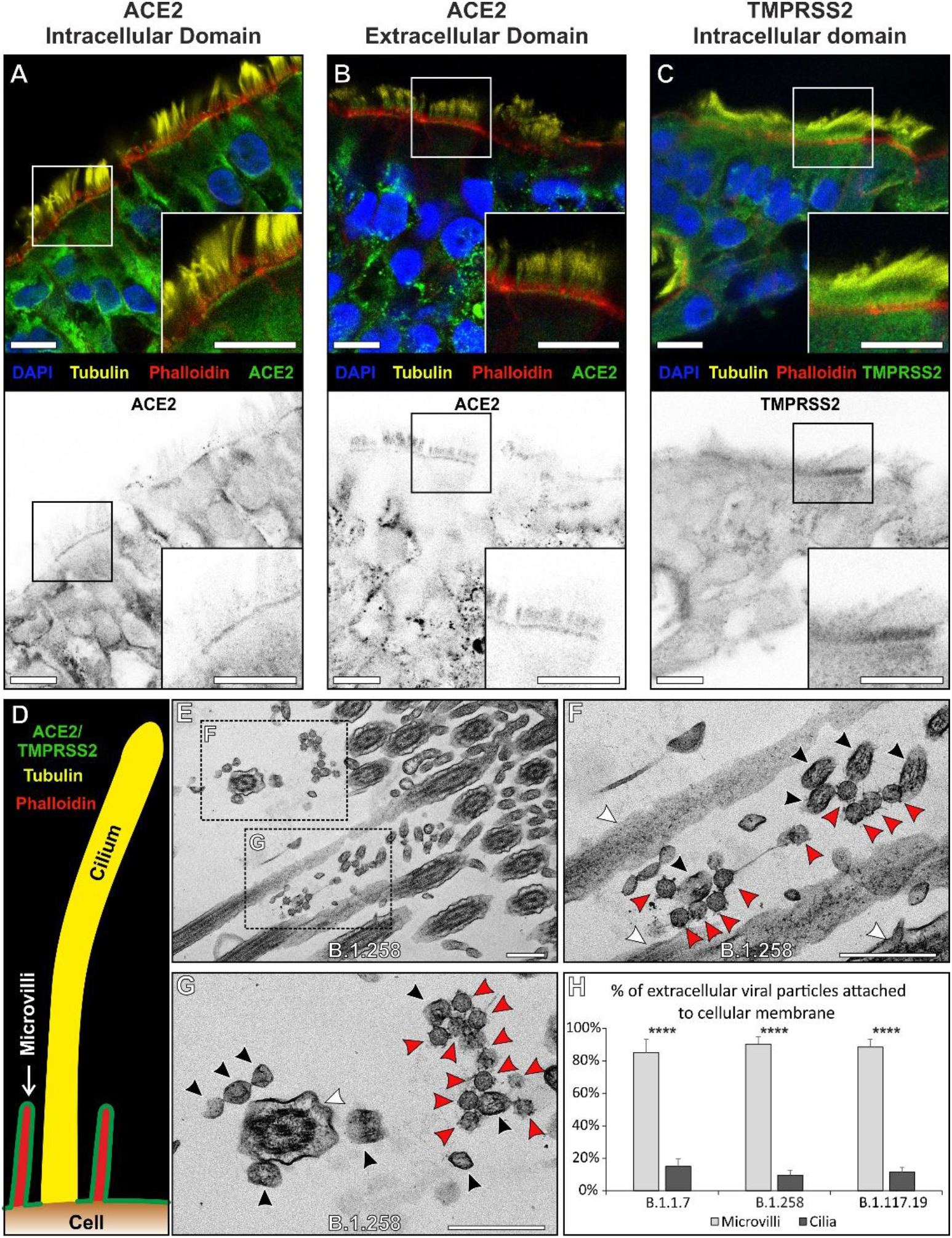
ACE2 and TMPRSS2 localises to the plasma membrane excluding cilia and this coincides with the sites of viral attachment. (A - C) Immunofluorescence antibody labelling against ACE2 and TMPRSS2 using anti-tubulin as a marker for cilia and phalloidin for the actin enriched microvilli. Both ACE2 and TMPRSS2 were found to be localised to cell surface that included the microvilli. Although there is diffuse, likely non-specific, staining everywhere with the TMPRSS2 antibody it is clearly enriched on the apical plasma membrane. Grey scale images of the staining profiles of ACE2 and TMPRSS2 show that they only extend a short difference from the cell surface unlike the tubulin staining of cilia and overlap with the phalloidin staining. (D) A diagram of the staining pattern of the antibodies. (E – G) When looking by TEM, virions (red arrowheads) are seen attached to microvilli (black arrowheads) and not to cilia (white arrowheads). (H) Quantification of the number of viral particles attached (within 3 nm) to microvilli and cilia, there is little affinity of the virus to cilia for all three isolates. Statistical significance was determine using Student’s t-tests (**** P ≤ 0.0001). Scalebars (A – C) 10 μm and (E – G) 400 nm

### Viral infection cannot be detected in goblet cells

VCs can be identified by TEM by the distinct morphology of virions contained within as shown in Figure 1 and Figure 4 A - C. VCs can be used as a marker of cellular infection as these will have resulted from successful viral entry and genome replication within the host cell. Few VC or VC-like compartments were identified when examining goblet cells that are characterised by the presence of mucin containing secretory granules (Mu in Figure 4 D). Endosome and endolyosomes (Figure 4 E) were observed in this cell type and could be distinguished from VCs by the intraluminal vesicles having a smaller diameter, less electron dense contents and lacking S glycoprotein (see Figure 2). The lack of VCs in goblet cells in comparison to ciliated cells was consistent for all isolates (Figure 4 F).

**Figure 4.**
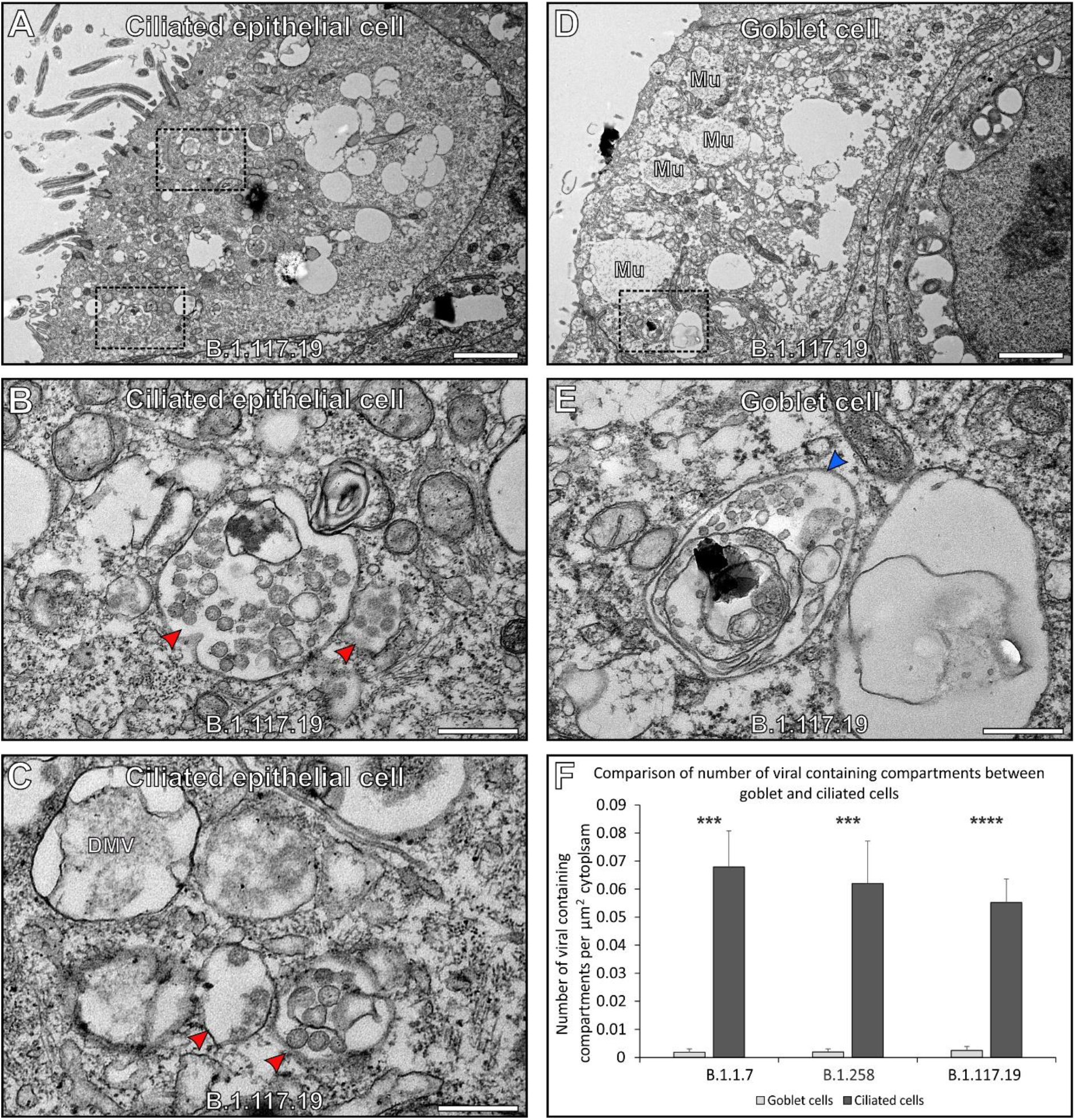
SARS-CoV-2 does not infect goblet cells. (A – C) Ciliated cells contain readily identifiable viral containing compartments (red arrowhead). (D, E) Goblet cells contain endosome and endolysosomes (blue arrowhead) that have a similar appearance to viral containing compartments but can be differentiated by the lack of viral nucleocapsid protein and S-glycoprotein structures. Goblet cells also contain mucin containing secretory granules (Mu). (F) Very few VCs were found in goblet cells for all three isolates. Statistical significance was determine using Student’s t-tests (*** P ≤ 0.001 and **** P ≤ 0.0001). Scalebars (A & D) 2 μm and (B, C & E) 400 nm.

### S glycoprotein like structures on the plasma of ciliated cells are a marker of infected cells

Another marker of infected cells was the presence of S glycoprotein protein-like structures at the plasma membrane. These were found on microvilli of infected ciliated cells but were absent from the microvilli of uninfected goblet cells (Figure 5). This feature can be seen when comparing a ciliated cell neighbouring a goblet cell (Figure 5 A – C). The goblet cell has microvilli with a smooth plasma membrane similar to that of an uninfected ciliated cell (Figure 5 B, D). S glycoprotein like protrusions are clearly visible on the infected ciliated cells microvilli (blue arrows in Figure 5 C and Supplementary Figure 5 A). Protrusions were absent from cilia of infected cells as shown in Supplementary Figure 5 B. Areas of the plasma membrane that have attached virions, also have protrusions with a similar morphology to the viral S glycoprotein (blue and red arrows in Figure 5 E-G). This could be a result of virus fusing with the plasma membrane and leaving behind S glycoprotein in the fused membrane patch, or S glycoprotein that has been expressed in an infected cell and transported to the plasma membrane. When examining cells from a nasal brushing of a SARS-CoV-2 infected patient (unknown viral lineage), similar protrusions on the plasma membrane and microvilli could be seen (Figure 5 H – J). In non-infected patients these protrusions were absent as seen in Supplementary Figure 5 C.

**Figure 5.**
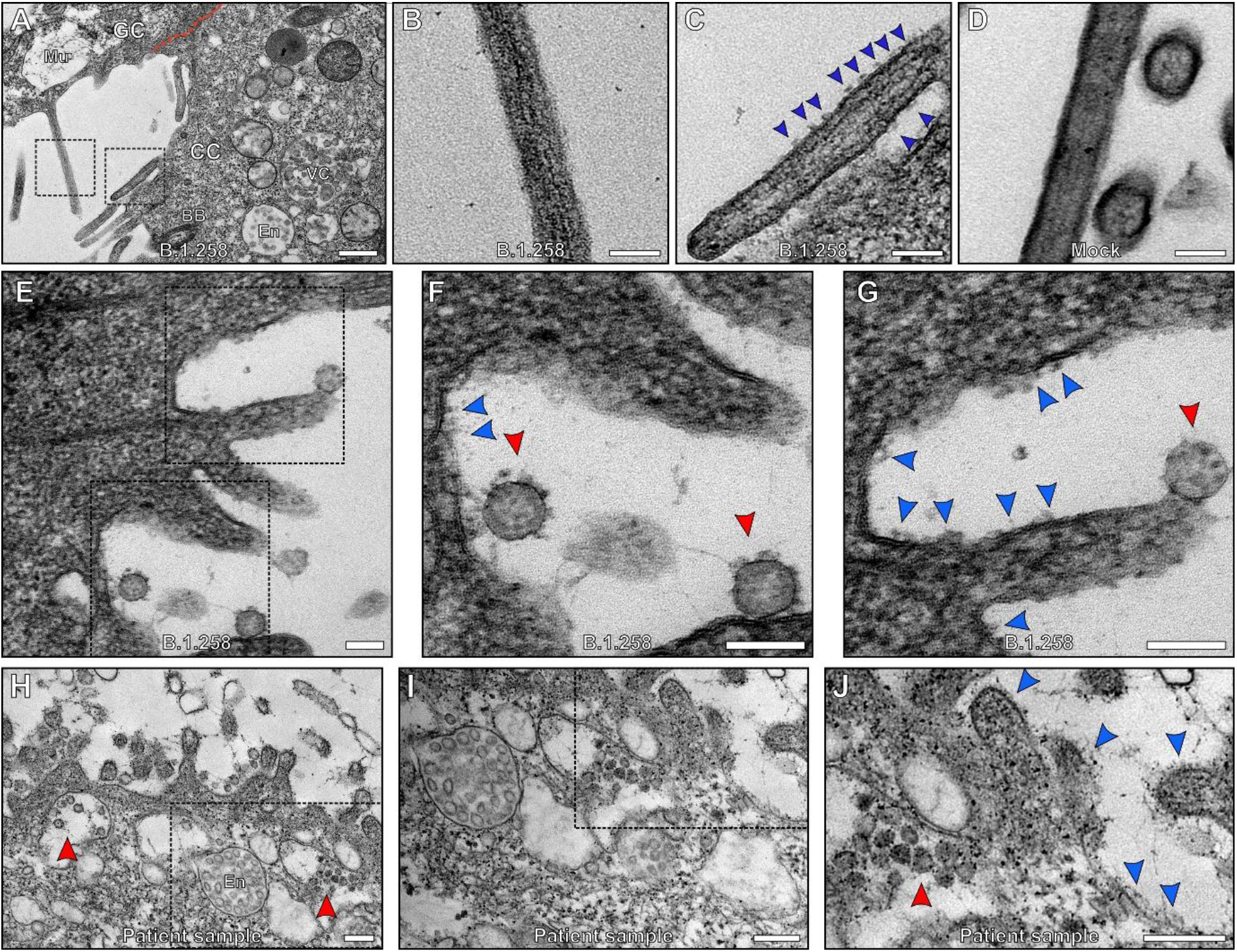
S-glycoprotein like protrusions were found on microvilli and plasma membrane of infected cells. (A) An uninfected goblet cell (GC) next to an infected ciliated cell (CC) that has viral containing compartments (VC) (endosome (En) were also visible). The goblet cell is identifiable by the mucin containing secretory granules (Mu) and the junction between the two cells false coloured with a red line. When looking at the microvilli from the goblet cell shown in (A) and at higher magnification (B), the membrane is smooth. Conversely, the plasma membrane of the microvilli of the infected ciliated cell (A) and at higher magnification (C), is not smooth and has protusions (blue arrowheads). (D) The microvilli of a non-infected ciliated cells are smooth. (E) and high magnification images of the boxed regions (F & G) show infected cultured human airway epithelial cells where viral particles can be seen attached to the cell surface. The viral S-glycoprotein (red) has a similar profile to the cell surface protrusions (blue). (H – J) Cells acquired from a nasal brushing of an infected patient show the presence of viral particles (red arrowheads). (I – J) At higher magnification protusions are seen at the cell surface and on microvilli (blue arrowheads). Scalebars (A) 400 nm, (B – G) 100 nm and (H – J) 200 nm.

### Viral fusion at the plasma membrane

A virion fusing at the host ciliated cell plasma membrane was captured by TEM and electron tomography (Figure 6). The membrane of a virion can be seen distended and appears fused to the host cell plasma membrane. Other virions are seen in the same vicinity (Figure 6 A, B). To examine the fusion in greater detail a tomogram was generated that shows the membrane of the virus is continuous with the plasma membrane of the host cell (Figure 6 C, D and shown in red in E, F). The nucleocapsid protein content within this virion is diluted compared to that of whole virions in the vicinity (false coloured in blue in Figure 6 E, F), indicating that some has already entered the host cell, consistent with this profile indicating viral fusion rather than budding. A neck-like structure was observed at the site of fusion, false coloured in yellow in Figure 6 E, F.

**Figure 6.**
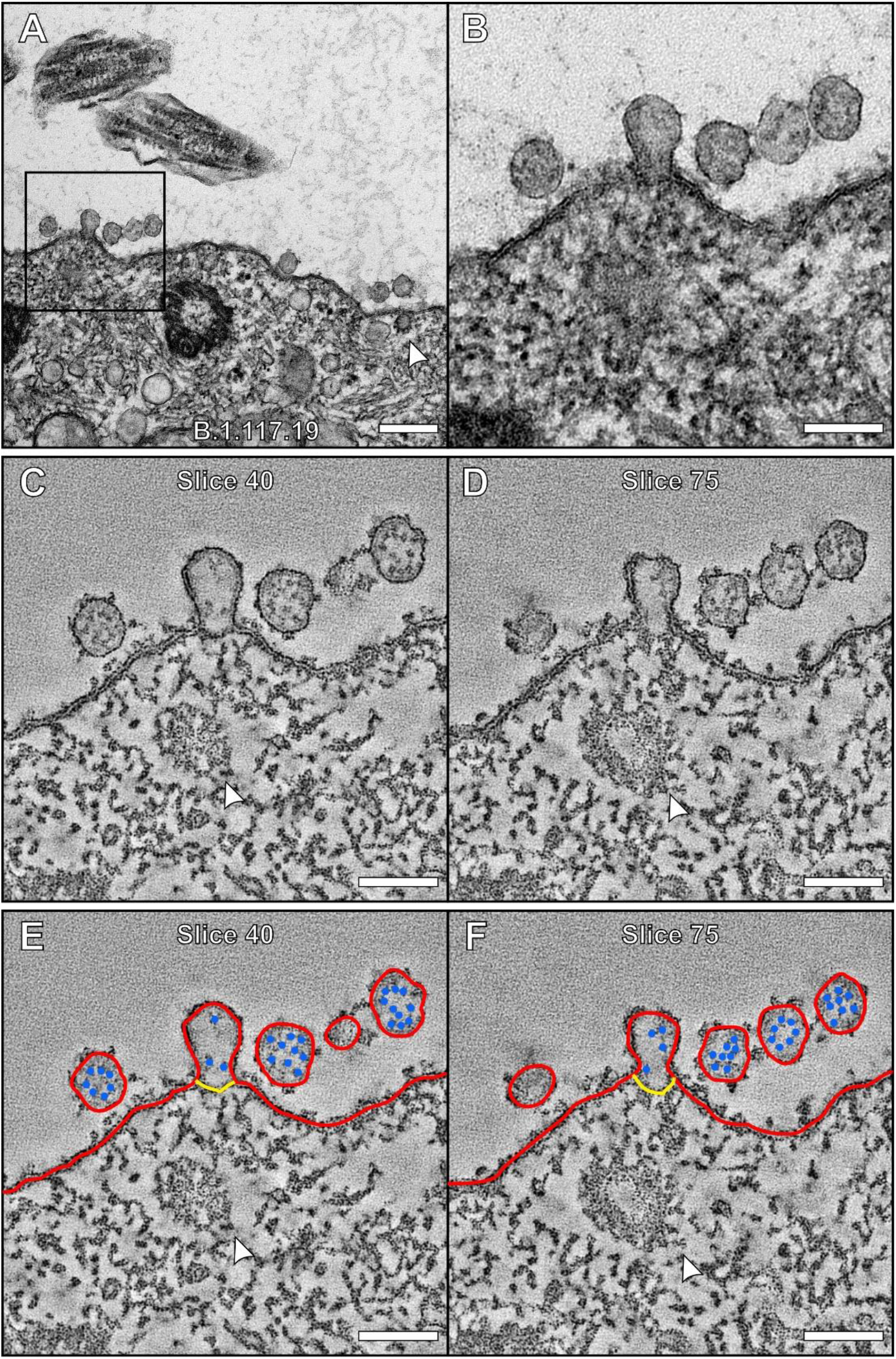
SARS-CoV-2 virion fused to the plasma membrane and releasing its contents into a ciliated cell. (A)TEM image of a virion fused to a host cell and higher magnification of the boxed region (B). (C – F) Tomogram generated of the attached virion with false colours (E, F) to indicate viral and plasma membrane (red), nucleocapsid protein (blue) and a neck-like structure (yellow). (E and F) The viral membrane is continuous with the host cell plasma membrane and the nucleocapsid contents diluted as they are released into the host cell. The white arrowheads highlight (A, B) a clathrin coated pit and (E, F) a clathrin coated vesicle. Scalebars (A) 200nm and (B – F) 100nm

### Viral budding within infected cells

Within VCs, virions were observed budding from the membrane (Figure 7 A, B). The neck-like structure at the site of viral attachment to the membrane indicates that the virion is pulling away in the direction of the VC lumen as shown in Figure 7 A. This implies budding rather than a virion fusing with the membranes. As these virions emerge from the membrane, they are clearly coated with S glycoprotein (Figure 7 A, B). A tomogram of a VC containing two budding virions shows a network of microtubules running into or near the emerging virions (yellow arrowhead in Figure 7 C, D). These could have a role in transporting viral components such a nucleocapsid protein and RNA to the site of viral budding. The electron dense nucleocapsid protein appeared to be concentrated at the periphery of the budding virions against the viral membrane, potentially driving the budding process (green arrowhead in Figure 7 C, D). This appearance of the nucleocapsid protein is different from that of the virion fused at the plasma membrane where it was found to be diluted (Figure 6), providing further evidence that these are budding profiles within VCs. Within DMVs and VCs, S glycoprotein like protrusions facing the lumen were observed, similar to the structures found on the plasma membrane of infected cells (Figure 7 E, F). To examine these structures further, tomograms were generated (Figure 7 G, H). These showed the structure of these protrusions in greater detail and confirmed they are likely to be S glycoprotein. Virions were also visualised within irregular shaped membrane structures in close contact to the VCs (red arrowheads) that appear to be convoluted membranes (Snijder et al., 2020).

**Figure 7.**
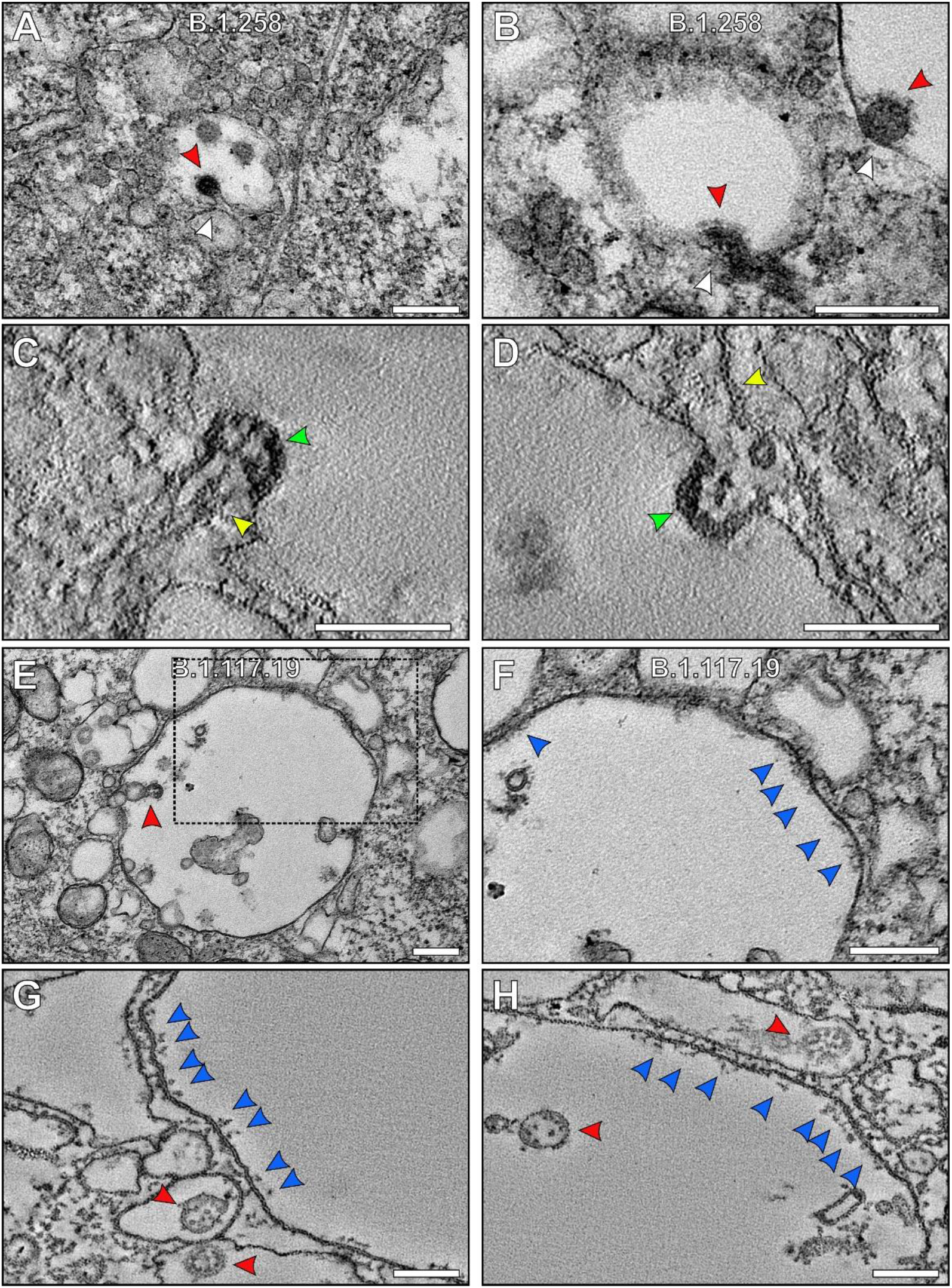
Virions bud into VCs coated in S-glycoprotein that are transferred from the limiting membrane. (A & B) Virions (red arrowheads) seen budding into a VC which has a S-glycoprotein coat. (C & D) Slices from a tomogram showing budding at the limiting membrane of a VC. Microtubules were seen running into budding virions (C) or nearby (D) as indicated by the yellow arrowheads. The nucleocapsid protein appeared to be concentrated at the membrane of the budding virions (green arrowhead). The boxed region in (E) shown at higher magnification (F) of an VC containing virus (red arrowhead) has S-glycoprotein (blue arrowheads) on the limiting membrane facing the lumen. (G & H) Slices from tomograms generated of VCs showing S-glycoprotein on the limiting membrane in more detail. Viral particles were within the VCS and DMVs as well as surrounded by membrane in structures close by. Scalebars (A, B, E, F) 200 nm and (C, D, G, H) 100 nm.

### Structural comparison of S glycoprotein and membrane protrusions

To further assess the structures of the protrusions at the plasma membrane and within VCs and to compared to viral S glycoprotein, subtomographic averages were generated. This approach removes background and enhance the 3D resolution of the tomography data as shown in Figure 8. All three reconstructions had a strong resemblance that included a globular-like structure protruding from either the viral, plasma or VC membrane. We also transfected HeLa cells with ss-HA-S glycoprotein and again observed protrusions at the plasma membrane (supplementary figure 6). Based on the 3D structures and presence of protrusions in ss-HA-S glycoprotein expressing HeLa cells it is highly likely that these are all S glycoprotein.

**Figure 8.**
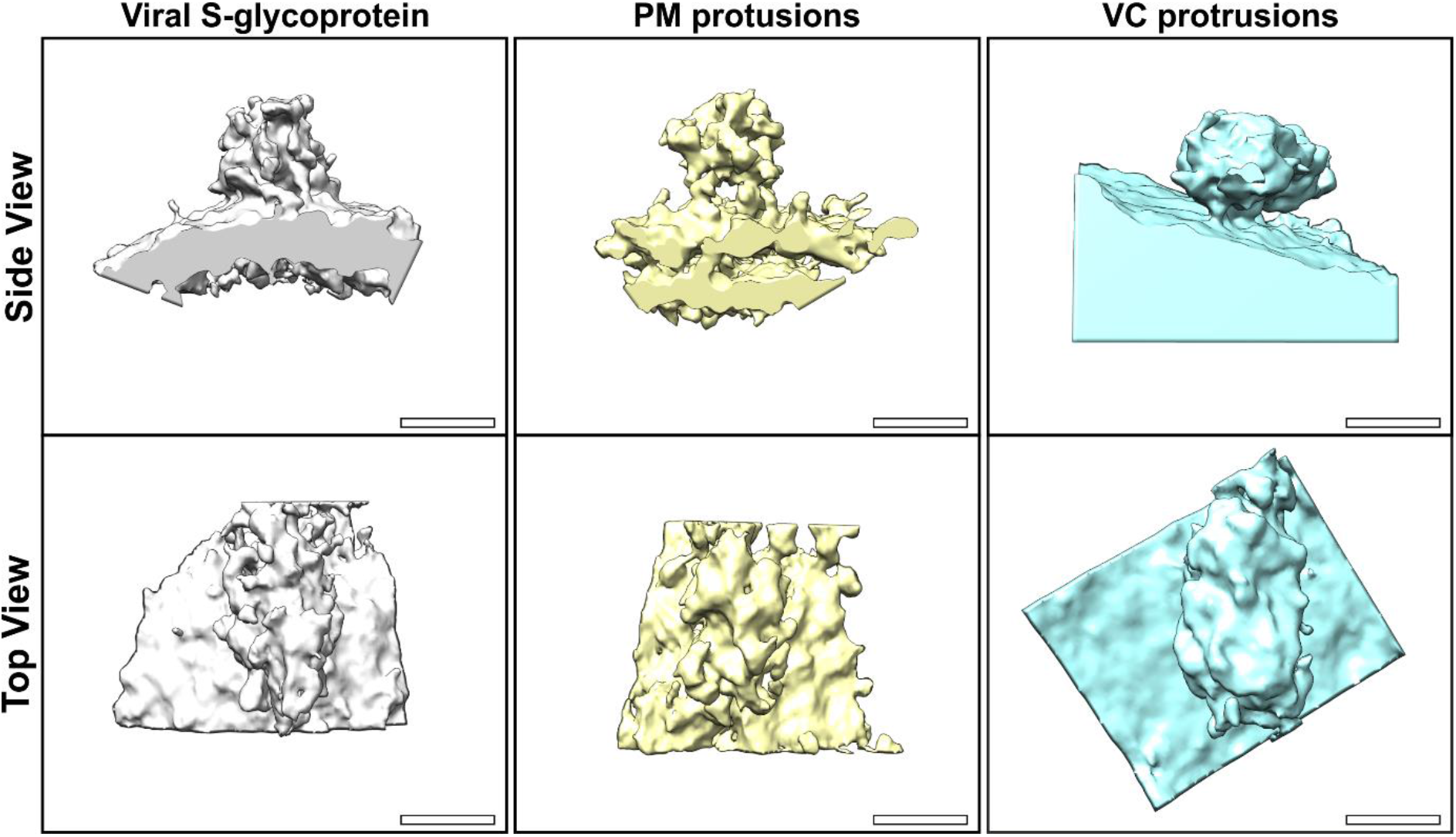
Subtomographic averages of plasma membrane (PM) and viral compartment (VC) protrusion have a similar structure to viral S-glycoprotein. Scalebar 10nm.

## Discussion

Electron microscopy techniques allow high magnification and detailed visualisation of viral processes that cannot be obtained using other methods. By examining primary human airway epithelial (HAE) cells infected with three different isolates of SARS-CoV-2 we are able to show aspects of viral attachment, entry and budding that have not been previously described. Due to the severity and global impact of SARS-CoV-2 it is essential to understand these processes as the mechanism underlying them present potential therapeutic targets.

For both SARS CoV and SARS CoV-2, ACE2 is a key host molecule expressed in airway cells required for viral entry through interaction with S glycoprotein (Gallagher and Buchmeier, 2001; Simmons et al., 2013; Hoffmann et al., 2020). In addition to engagement with the ACE2 receptor, SARS CoV2 utilizes a second host molecule, TMPRSS2, that cleaves the S glycoprotein S2 subunit to allow membrane fusion and release of viral contents into the host cell. Previous studies have indicated ACE2 is localised to the cilia plasma membrane, which implies that cilia would be the likely site of extracellular virion attachment (Lee et al., 2020). In contrast, we show here both ACE2 and TMPRSS2 are localised to discrete areas of the ciliated cell plasma membrane that include microvilli but excludes cilia themselves. A number of factors could underlie this discrepancy, including the degree of differentiation of the cells and the sample preparation, given that the previous study used paraffin embedded samples that require antigen retrieval unlike the cryostat sections examined here. Our conclusion that ACE2 is largely excluded from the cilia is strengthened by the ease of distinction between cilia and microvilli in our highly differentiated cells, our demonstration that these antibodies faithfully localise transfected ACE2 in cells that do not normally express it and that the virions attach preferentially to microvilli and other regions of the plasma membrane and rarely associate with cilia

Goblet epithelial cells that neighbour ciliated epithelial cells have microvilli but lack cilia. When comparing these cell types in infected HAEs, evidence of intracellular virus replication such as VCs could be clearly seen in ciliated cells but were absent from goblet cells. Similar observations were made in a previous study looking at a seasonal coronavirus (Afzelius, 1994; Dijkman et al., 2013). It is important to note that no DMVs were observed within goblet cells, and virions were not seen at the cell surface (data not shown). This shows that goblet cells likely do not possess the required machinery for viral entry either via an extracellular route or laterally from neighbouring ciliated cells. Over the 72 hours of SARS-CoV-2 infection, there was extensive viral replication as indicated by the large number of extracellular virions exclusively at the surface of ciliated cells and measurement of replication kinetics in a parallel study (Brown et al., 2021). Due to this large viral load, we predict it is unlikely with an extended time of infection that the goblet cells would become vulnerable to infection, but this would need to be tested in a further study.

Viral fusion at the plasma membrane is proposed as a route of viral entry for SARS-CoV-2 in airway cells (Peacock et al., 2020). We were able to capture and examine this process using the 3D technique, electron tomography. The viral membrane could be seen to be continuous with the host cell plasma membrane and its contents diluted as they likely flowed from the viral lumen into the host cell cytoplasm. A neck-like structure was seen at the site of membrane fusion that may indicate that the virus has only just fused or there is something regulating the shape of the membranes at the fusion site. After virion fusion and release of their contents, these structures must flatten otherwise a large number of fused viral structures would be seen on the plasma membrane of infected cells. As this occurs S glycoprotein that was integrated into the viral membrane would remain at the plasma membrane. We were able to visualise the presence of S glycoprotein like structures on the surface of infected cells that could have resulted from previous viral fusion processes.

Viral budding was visualised as newly forming virions emerging from the membrane of VCs. Due to the profile of budding events where the bud appeared to be pulling away from the membrane it is unlikely that these are fusion events. The VCs appear spherical in shape when examining EM sections and some were seen to position close to recognisable Golgi-like membrane but is difficult to be certain if they are ERGIC derived structures based on their morphology alone. Further work such as immuno-electron microscopy could help to determine the proteins localised to the VCs that have budding virions. In close proximity to the VCs, virions were found in less regular shaped membrane structures that are likely to be convoluted membranes. More work is required to determine the origin of the VCs and how virions are able to form in these proximal membranous structures. At the membrane of VCs S glycoprotein like structures were seen facing the lumen. As virions form and bud this would allow them to gain their envelope containing S glycoprotein. If the processes that leads to expression and/or transport of S glycoprotein are not always optimal this could explain why some virions were found within VCs without a S glycoprotein coat. If virions are released from host cells via fusion of VCs with the plasma membrane, any remaining S glycoprotein on the membrane would likely be transferred to the cell plasma membrane and would face the extracellular space. This could also be the source of the S glycoprotein detected at the surface of infected cells and could results in cell-cell fusion that has been reported elsewhere by us and others (Braga et al., 2021, 16). Evidence was recently presented that SARS CoV2 virions leave the cell via lysosome exocytosis (Ghosh et al., 2020). Although some of the VCs have membranous inclusions, most do not have content typically associated with lysosomes. Further work is needed to determine the nature of these compartments and their relationship with both ERGIC and the lysosome.

The presence of S glycoprotein on the plasma membrane and within the lumen of cellular compartments provide an excellent marker of infected cells by electron microscopy. Based on the subtomographic averages and the presence of protrusions by EM localised to the plasma membrane and within VCs after infection it is highly likely that they are S glycoprotein. More work is required to determine if cell surface S glycoprotein is a result of either viral fusion with incoming particles and/or release from VCs. No difference between the isolates was detected, as higher resolution methods such as cryo-electron microscopy would be required to identify molecular differences between the S glycoproteins that only differ by 9 or so amino acids. Even so, the similarities of the isolates provided an excellent overview of SARS-CoV-2 infection. There is still much more to be learned from ultrastructural analysis including the route to viral release and alternative routes of viral entry.

This study has allowed better characterisation of key processes in the life cycle of SARS-CoV-2. Cilia were found not to be the site of key entry molecules ACE2 and TMPRSS2 and hence not involved in viral attachment and entry. In contrast the microvilli on ciliated cells expressed abundant receptors and were a site for virion attachment followed by cell surface fusion. Goblet cells were refractory to infection. This knowledge as well as what can be gained in further studies to establish factors regulating viral budding has potential to help in the design of novel therapeutic interventions.

## Acknowledgments

This work was supported by Wellcome Trust Grants 093445 (to C.E.F.) and the G2P-UK National Virology Consortium funded by UKRI (to J.C.B and W.S.B).

