## Supplementary figures for "Ultrastructural insight into SARS-CoV-2 attachment, entry and budding in human airway epithelium"

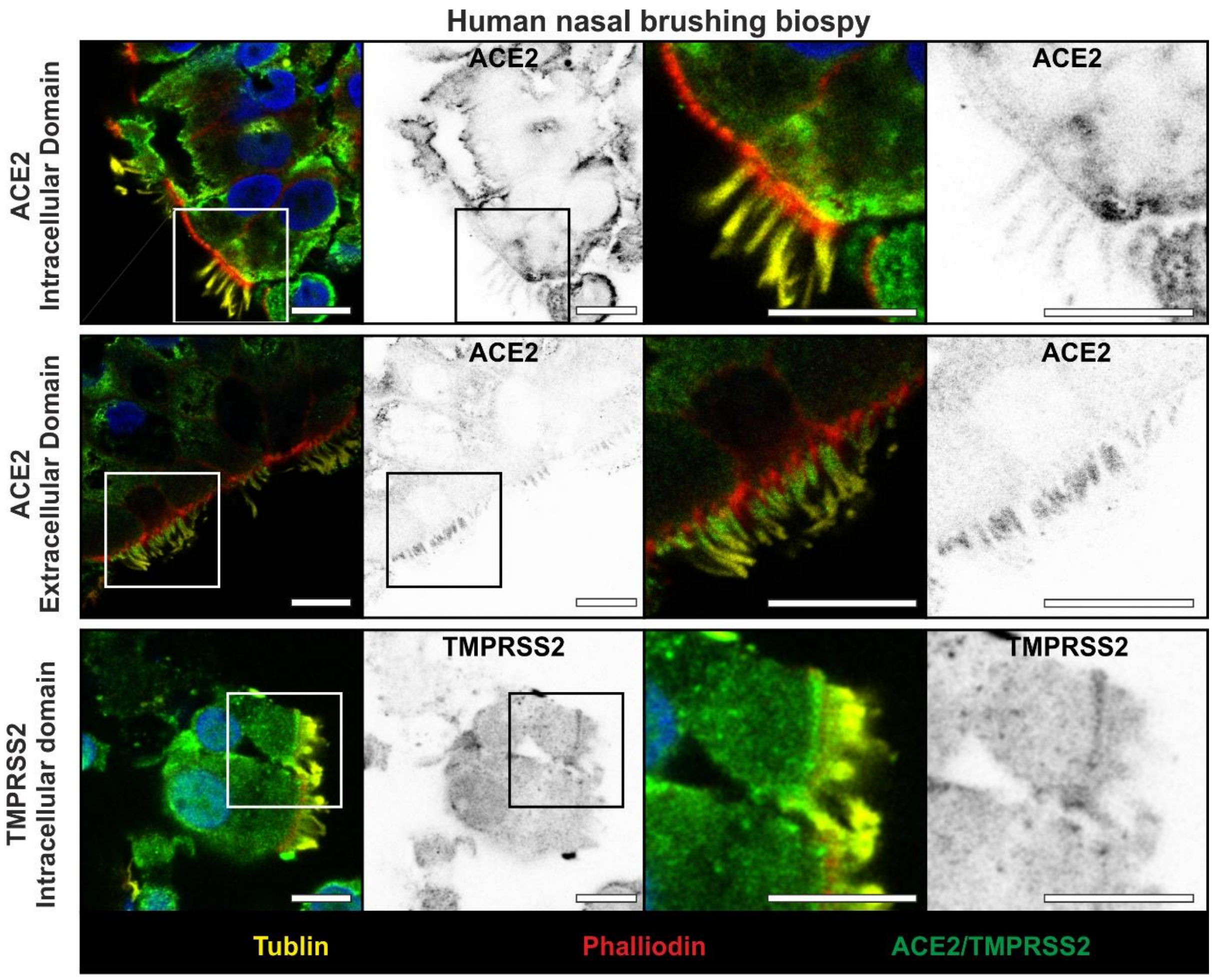


Supplementary figure 1. ACE2 and TMPRSS2 labelling of human nasal biopsy shows the same labelling pattern as cultured HAE. Scalebar 10um

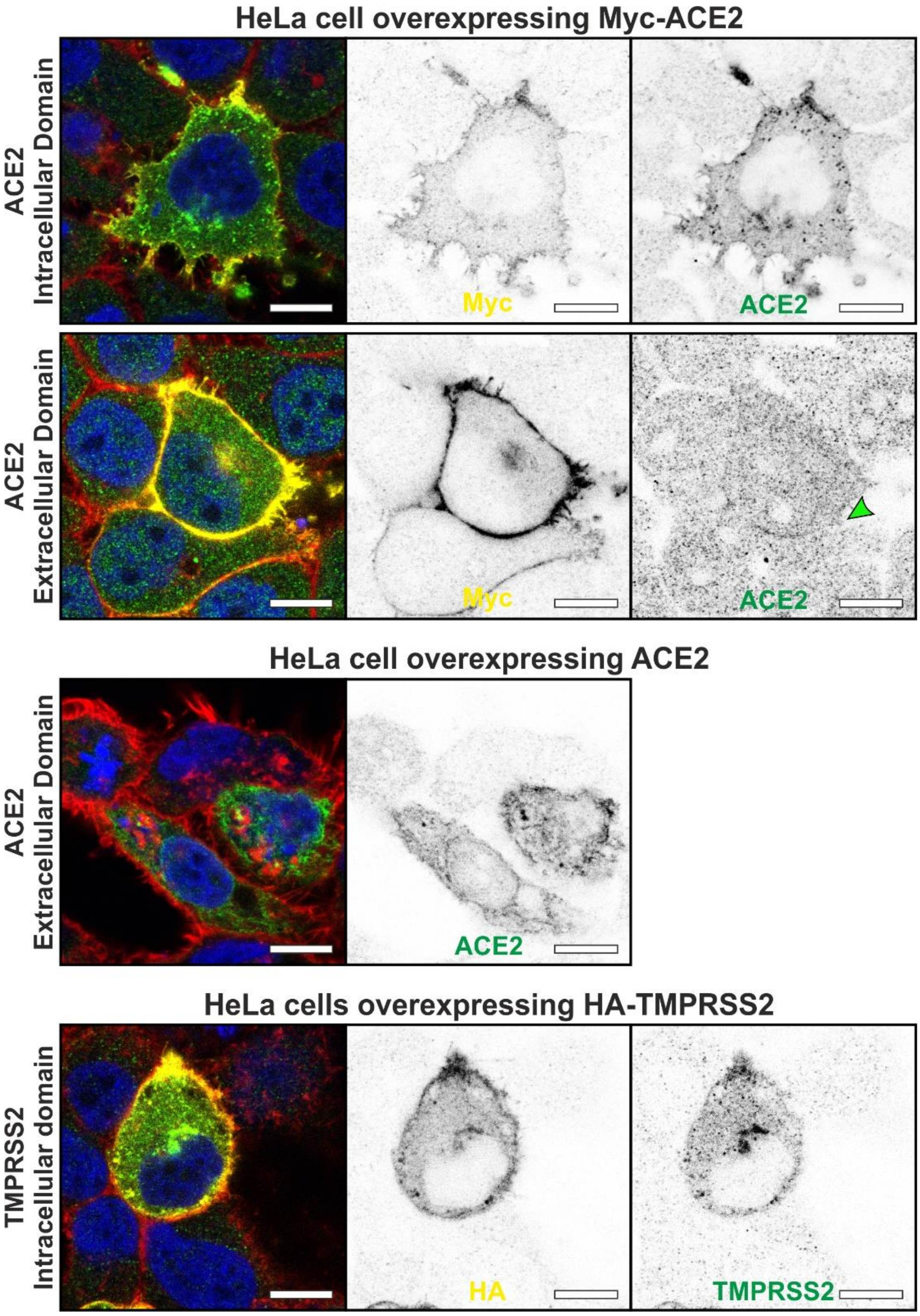


Supplementary figure 2. Validation of ACE2 and TMPRSS2 antibodies in HeLa cells expressing constructs. Signal from antibody staining overlapped with the labelling of Myc or HA in HeLa cell over expressing Myc-ACE2 or HA-TMPSS2. For Myc-ACE2, the Myc is an N-terminal tag which is the region the extracellular domain antibody is raised against, therefore, better antibody labelling is found in HeLa cells expressing untagged ACE2. Phalloidin staining is shown in red. Scalebar 10um

### ACE2 extracellular domain staining of non-permeabilised human respiratory epithelial section

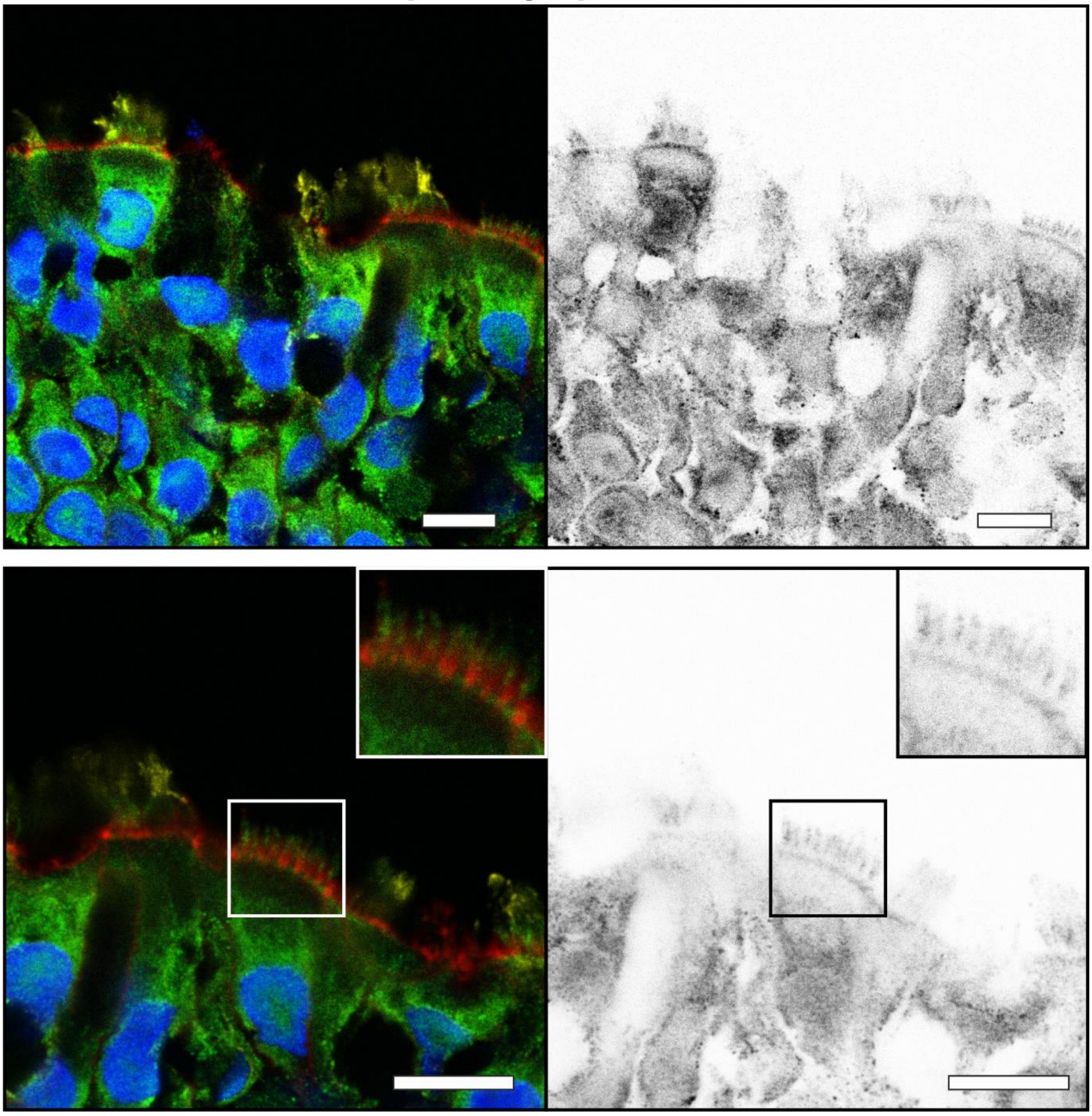

Supplementary figure 3. ACE2 antibody labelling against the extracellular domain of HAE that was not permeabilised showed localisation to microvilli and plasma membrane. The lack of permeabilised results in poor tubulin staining compared to permeabilised samples (see figure 2). The boxed region is an area of a ciliated cell in a slice where the cilia are absent and the ACE2 staining surround the actin (phalloidin stained)  enriched microvilli. Scalebar 10um

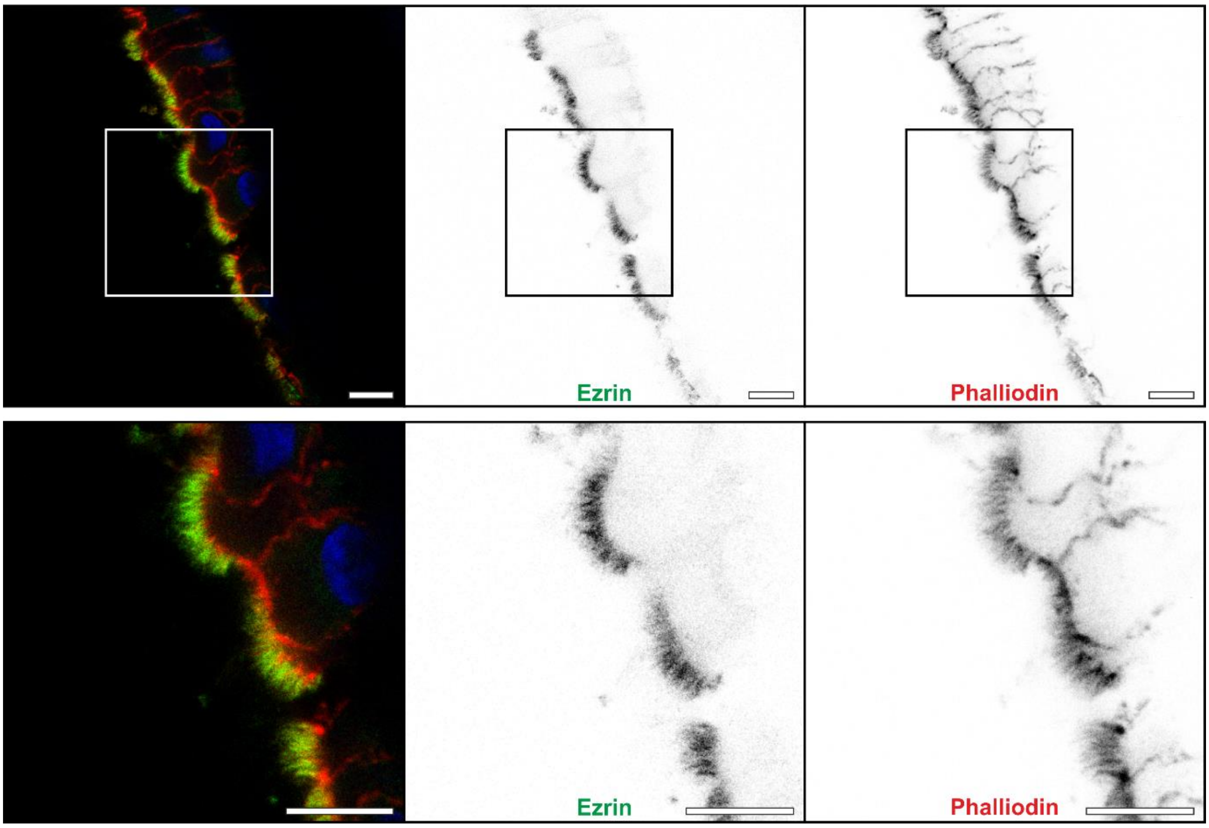


Supplementary figure 4. Ezrin localises to microvilli and the staining surround the actin (phalloidin stained). Boxed regions in the top panel are shown at higher magnification in the lower panels. Scale bar 10um.

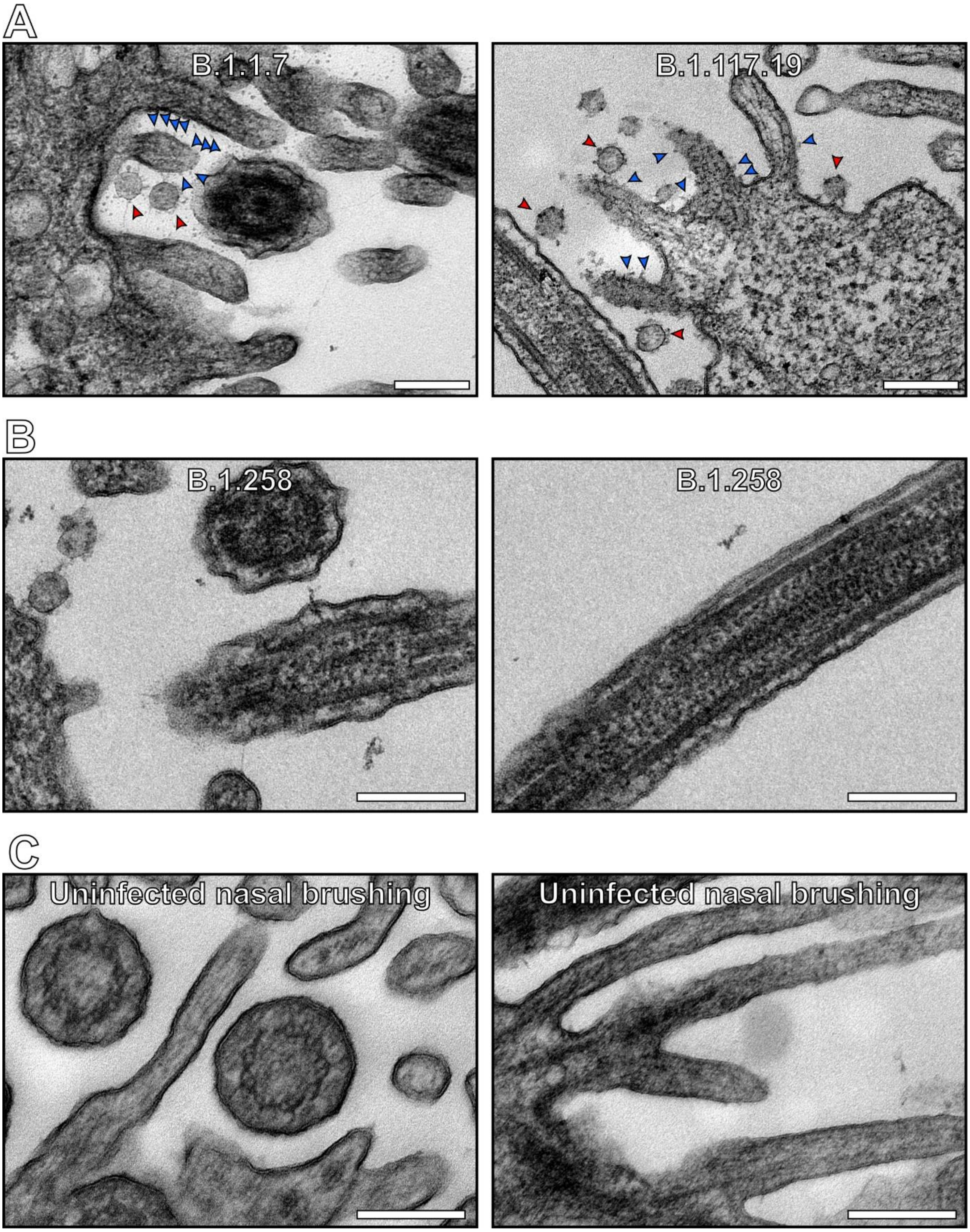


Supplementary figure 4. Infected ciliated cilia have protrusions on the plasma membrane apart from cilia. (A) Protrusions seen on the plasma membrane of microvilli (blue arrowheads) of infected (virions shown by red arrowheads) ciliated cells. (B) Ciliary membranes of an infected cell are smooth. (C) Microvilli and other regions of the plasma membrane are smooth in an uninfected nasal brushing sample. Scalebars 200nm

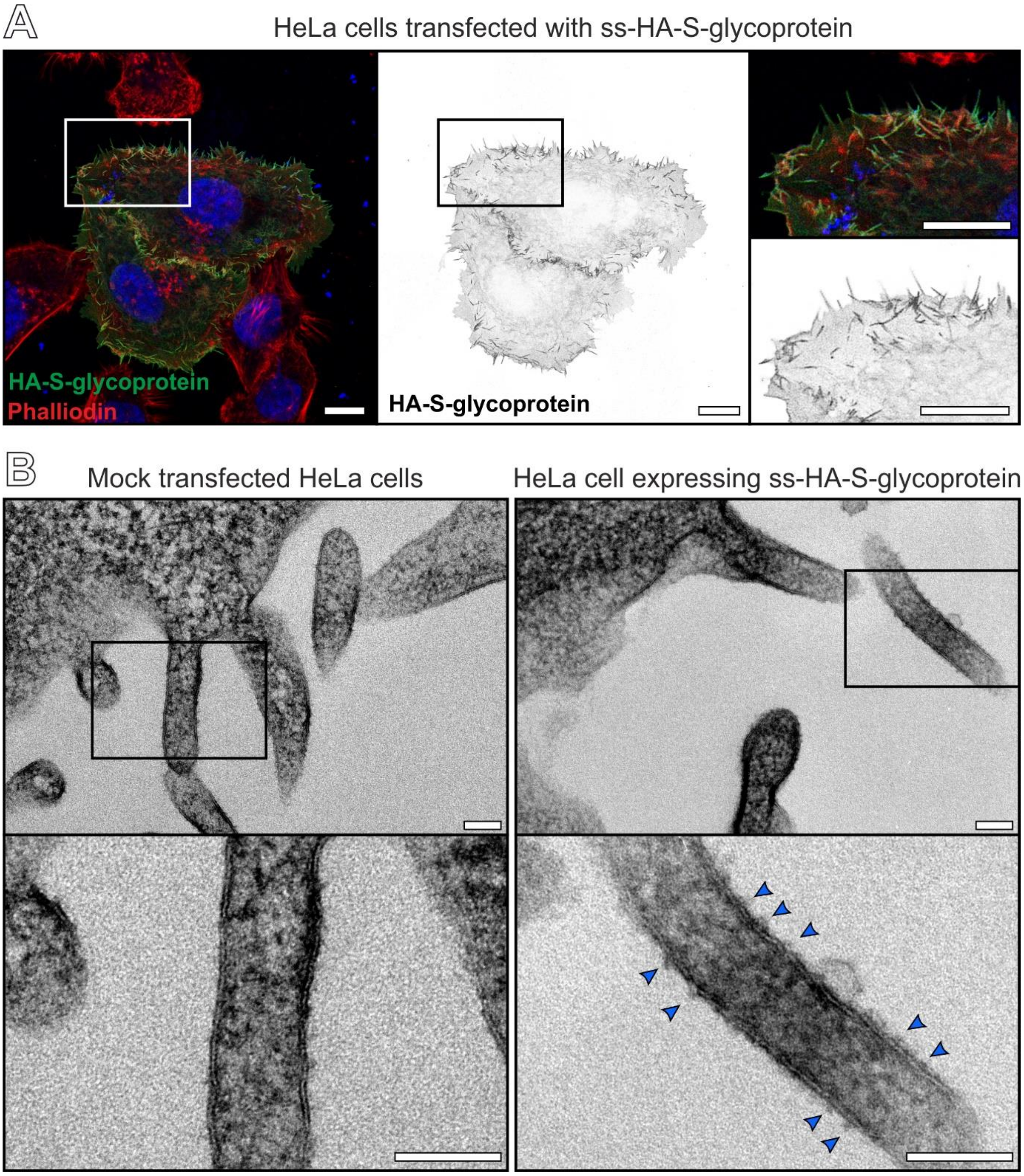


Supplementary figure 5. Overexpression of ss-HA-S-glycoprotein in HeLa cells leads to it being localised to the plasma membrane including to microvilli and results in protrusions as seen by EM. (A) IF images showing HA-S-glycoprotein localised to the plasma membrane in over expressing HeLa cells. HeLa cells over expressing ss-HA-S-glycoprotein have protrusion on the microvilli when compared mock infected cells. Scalebars (A) 10um and (B) 100nm.
